# Robustifying genomic classifiers to batch effects via ensemble learning

**DOI:** 10.1101/703587

**Authors:** Yuqing Zhang, W. Evan Johnson, Giovanni Parmigiani

## Abstract

Genomic data are often produced in batches due to practical restrictions, which may lead to unwanted variation in data caused by discrepancies across processing batches. Such “batch effects” often have negative impact on downstream biological analysis and need careful consideration. In practice, batch effects are usually addressed by specifically designed software, which merge the data from different batches, then estimate batch effects and remove them from the data. Here we focus on classification and prediction problems, and propose a different strategy based on ensemble learning. We first develop prediction models within each batch, then integrate them through ensemble weighting methods. In contrast to the typical approach of removing batch effects from the merged data, our method integrates predictions rather than data. We provide a systematic comparison between these two strategies, using studies targeting diverse populations infected with tuberculosis. In one study, we simulated increasing levels of heterogeneity across random subsets of the study, which we treat as simulated batches. We then use the two methods to develop a genomic classifier for the binary indicator of disease status. We evaluate the accuracy of prediction in another independent study targeting a different population cohort. We observed a turning point in the level of heterogeneity, after which our strategy of integrating predictions yields better discrimination in independent validation than the traditional method of integrating the data. These observations provide practical guidelines for handling batch effects in the development and evaluation of genomic classifiers.

## Introduction

Statistical learning models based on genomic information have been widely used for prognostication and prediction across a range of precision medicine applications, including cancer (Golub *et al.* (1999); Riester *et al.* (2014); Silvestri *et al.* (2015); Papaemmanuil *et al.* (2016)) and infectious diseases (Seib *et al.* (2009); Leong *et al.* (2018)), and have shown great potential in facilitating clinical and preventative decision making (Badani *et al.* (2015)). To fully achieve such potential, it is critical to develop prediction algorithms with generalizable prediction performance on independent data, which are transferable to clinical use (Simon *et al.* (2003)). However, the presence of “study effects”, or heterogeneity across genomic studies, makes it challenging to develop generalizable prediction models. In particular, it has been established that cross-study validation performance of genomic classifiers is often inferior to internal cross-validation (Ma *et al.* (2014); Chang and Geman (2015); Bernau *et al.* (2014)), and that this gap cannot be entirely explained by the most easily identifiable sources of study heterogeneity (Zhang *et al.* (2018)). Further research is needed to better understand and address the impact of heterogeneity on predictor performances.

In this study, we focus on a particular component of study effects, known as batch effects (Leek *et al.* (2010)), and aim to address its unwanted impact in binary classification problems. Batch effects are variation across batches of data due to differences in technical factors. Existence of batch effects threatens the reproducibility of genomic findings (Kupfer *et al.* (2012)). And therefore, it is necessary to develop efficient methods to remove the unfavorable influence of batch effects.

Many batch effect adjustment methods have been proposed for gene expression microarray (Leek and Storey (2007); Johnson *et al.* (2007); Gagnon-Bartsch and Speed (2012); Gagnon-Bartsch *et al.* (2013); Benito *et al.* (2004)) and sequencing data (Leek (2014); Risso *et al.* (2014)). These methods share the general idea to first merge all batches, estimate parameters representing differences in batch distributions, and then remove them from the data, resulting in a single adjusted dataset for downstream analysis. Here we adopt a different perspective, and propose to address batch effects with ensemble learning. Contrary to the traditional batch effect adjustment methods, our proposed framework is based on the integration of predictions rather than that of data. This is a simpler task for prediction, as it operates in one dimension rather than many. Also, it is possible to reward predictors that show good performance across batches and thus altogether ignore, rather than trying to repair, features that are preferentially affected by batch effects.

Although ensemble learning is a well established method, it has only recently been discussed in the context of training replicable predictors. Patil and Parmigiani (2018) found that ensembles of learners trained on multiple studies generate predictions with more replicable accuracy. In their framework, a cross-study learner (CSL) is specified by three choices: a) a data subsetting strategy; b) a list of one or more single-study learners (SSLs), which can be any machine learning algorithm producing a prediction model using a single study; and c) a combination approach utilizing multiple prediction models to deliver a single prediction rule. In our case, we subset the data by batch, and use the same CSL with batches in place of studies. Guan *et al.* (2019) provide theoretical insights in the comparison between merging and ensembling in training learners from multiple studies, and conclude that although merging studies is better than ensembling when the studies are relatively homogeneous, ensembling yields better performing models when the level of study heterogeneity is higher.

In this paper, we explore using ensemble learning in the context of batch effect adjustment for the first time. We provide both realistic simulations and real data examples to demonstrate the utility of our ensembling framework, and compare it with traditional merging strategies for addressing batch effects.

## Materials and methods

### Addressing batch effects via ensemble learning

We structure the problem as follows: in a binary classification problem on genomic data, we have a training set for learning prediction models (*S_trn_*), and another independent test set (*S_tst_*) where predictions are to be made. Subjects in the training set are associated with a binary label indicating their phenotype. Examples of the phenotype could be disease status (e.g. cancer versus normal) or response to a treatment. In addition, expressions of genes for all individuals are profiled for use as predictors. Individuals in the test set also have measured gene expressions, and the goal is to train a model that can accurately predict the disease label for them based on their gene expression profiles. We assume that the training set is generated in *B* batches 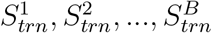 possibly due to practical or technical restrictions. Each batch contains a sufficient number of samples to train a prediction model. We assume both additive and multiplicative batch effects (Johnson *et al.* (2007)), which cause differences in the mean and the variance of gene expressions across batches.

Consider a collection of *L* learning algorithms to use for training. Multi-study learning begins by training each of the algorithms within each of the batches. This results in the collection 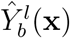, *l* = 1,…*L* and *b* = 1*,… B*, where 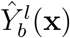 is the prediction function trained on batch *b* with learning algorithm *l*. In the binary classification setting, 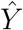 are the probabilities of samples belonging to the positive class. The final cross-study learner’s (CSL) prediction is calculated by a weighted average of predictions from each model, that is:

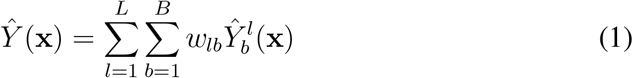

The performance of a CSL relies critically on the weights *w_lb_*, as these have the function of rewarding elements of the ensemble which show stable predictive performance across batches. We explore five weighting strategies, which fall into three categories, as described in Patil and Parmigiani (2018). The first is sample size weights, which uses scaled batch sizes as weights for models trained in the corresponding batches, and makes not effort to reward robustness to batches via weights. The second is cross-study weights, for which we evaluate how well each learned model performs when applied to the other batches within the training set, and assign higher weights to models that have better prediction performances. The last category is stacking regression weights (Breiman (1996)), for which we use each model to make predictions of the training data, and estimate the weights as regression coefficients between stacked predictions of the training samples and their labels. The association coefficients are estimated using non-negative least squares. A more detailed description of the weighting strategies is available in the Supplementary Materials.

### Data

We use a collection of RNA-Seq and microarray studies targeting subjects infected with tuberculosis (TB) to apply and evaluate our ensemble learning method, and make comparisons with the traditional strategy of batch adjustment followed by merging (for short “merging”). Subjects involved in this collection of studies can be divided in three phenotypes based on their disease progression status: 1) latent infection / non-progressing, 2) in “progression”, or those that will progress to disease in the near future, and 3) active TB disease. For simplicity and sample size considerations, we use two types of patients in each analysis to form a binary classification problem, and focus on different phenotypes for simulation and application of our methods. In simulations, we focus on predicting progressors against non-progressors using publically available studies, as this separation yields the largest sample size in the training set. In real data, since there are no known batch effects within any single study, we aim to separate subjects with latent infection from those with active disease instead, so we have at least three studies in the collection. We merge two studies for training while treating the differences between studies as “batch effects”. Table 1 summarizes information on the samples used in simulation studies and real data analysis.

**Table 1:**
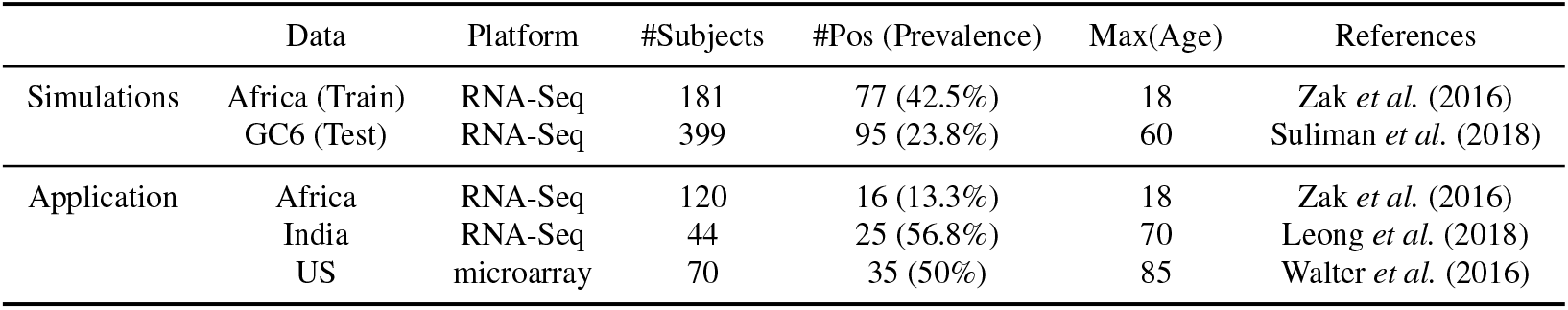
Summary statistics of the TB datasets used in simulation studies and real data applications. #Subjects: total number of individuals in each study. #Pos: number of positive samples. They refer to progressors in simulation studies, and active patients in real data application. The negative samples are subjects with latent infections in both cases. Prevalence: percentage of positive samples in the study. Max(Age): the maximum of age among all subjects enrolled in the studies. Note that the Africa study targets only adolescents between ages of 12 and 18, while the remaining studies have a wider range of age. The US study is generated from Affymetrix Human Gene 1.1 ST Array, while the rest are RNA-Seq studies.

|  | Data | Platform | #Subjects | #Pos (Prevalence) | Max(Age) | References |
| --- | --- | --- | --- | --- | --- | --- |
| Simulations | Africa (Train) | RNA-Seq | 181 | 77 (42.5%) | 18 | Zak *et al.* (2016) |
|  | GC6 (Test) | RNA-Seq | 399 | 95 (23.8%) | 60 | Suliman *et al.* (2018) |
| Application | Africa | RNA-Seq | 120 | 16 (13.3%) | 18 | Zak *et al.* (2016) |
|  | India | RNA-Seq | 44 | 25 (56.8%) | 70 | Leong *et al.* (2018) |
|  | US | microarray | 70 | 35 (50%) | 85 | Walter *et al.* (2016) |

### Simulation for comparing merging and ensembling

We consider the two datasets collected from the African population reported by Zak *et al.* (2016) and Suliman *et al.* (2018), as described in Table 1. We use one of the two studies for training, and the other for prediction. For the training set only, we randomly assign individuals to disjoint subsets which will be simulated to be processing batches, and simulate differences in the moments of gene expression distributions across batches as described below. No batch effects are added to the validation set. We train predictors using both merging and ensembling on this dataset with simulated batch effects, then make predictions in the other independent study. We evaluate the two approaches using discrimination in the independent study. Details follow.

#### Simulation of batch effects

We transformed the sequencing data into logFPKMs, and selected the top 1000 genes with the highest variances for building the classifiers. Then, we randomly took subsets of individuals from the training set to form 3 batches, each batch containing 10 non-progressors and 10 progressors. We then simulated batch effects across the 3 batches.

Our data generating model for batch effects is the linear model assumed in the ComBat batch adjustment method (Johnson *et al.* (2007)). Specifically, we estimate two components from the original training data: 1) the expression of gene *g* among the negative samples, and 2) the biological effect (i.e. the expression changes due to biological perturbations or conditions of interest). We then specify batch effect parameters affecting the mean (*γ_gb_*) and the variance (*δ_gb_*) of expression in gene *g* caused by batch *b*. Same as in Johnson *et al.* (2007), *γ_gb_* and *δ_gb_* are randomly drawn from hyper-distributions

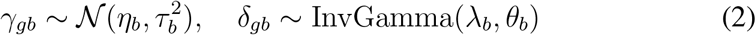

ComBat assumes an additive batch effect for the mean, and a multiplicative batch effect for the variance. To set the hyper-parameters, we first specify a value to represent the severity of batch effects, as reported in columns “Batch Effect on Mean” (*sev_mean_*) and “Batch Effect on Variance” (*sev_var_*) in Table 2. We selected 3 severity levels (*sev_mean_* ∈ {0, 3, 5}) for batch effect on the mean, and 5 levels (*sev_var_* ∈ {1, 2, 3, 4, 5}) for batch effect on variance. Given a severity level for batch effects, we fixed values for *τ_b_* s a nd *θ_b_* s, so that the variance of *γ*_*gb*_ and *δ_gb_* over genes are 0.01. We varied the mean of these two parameters, so that the hyper mean *η*_*b*_ is (*−sev_mean_,* 0, +*sev_mean_*), and the hyper variance *λ*_*b*_ is 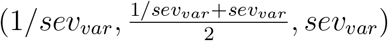 for the three batches. The parameters are then added or multiplied to the expression mean and variance of the original study. The characteristics of simulated batches are also summarized in Table 2. Figure 1 shows an example training set where we simulated 3 batches with both mean and variance differences.

**Table 2:**
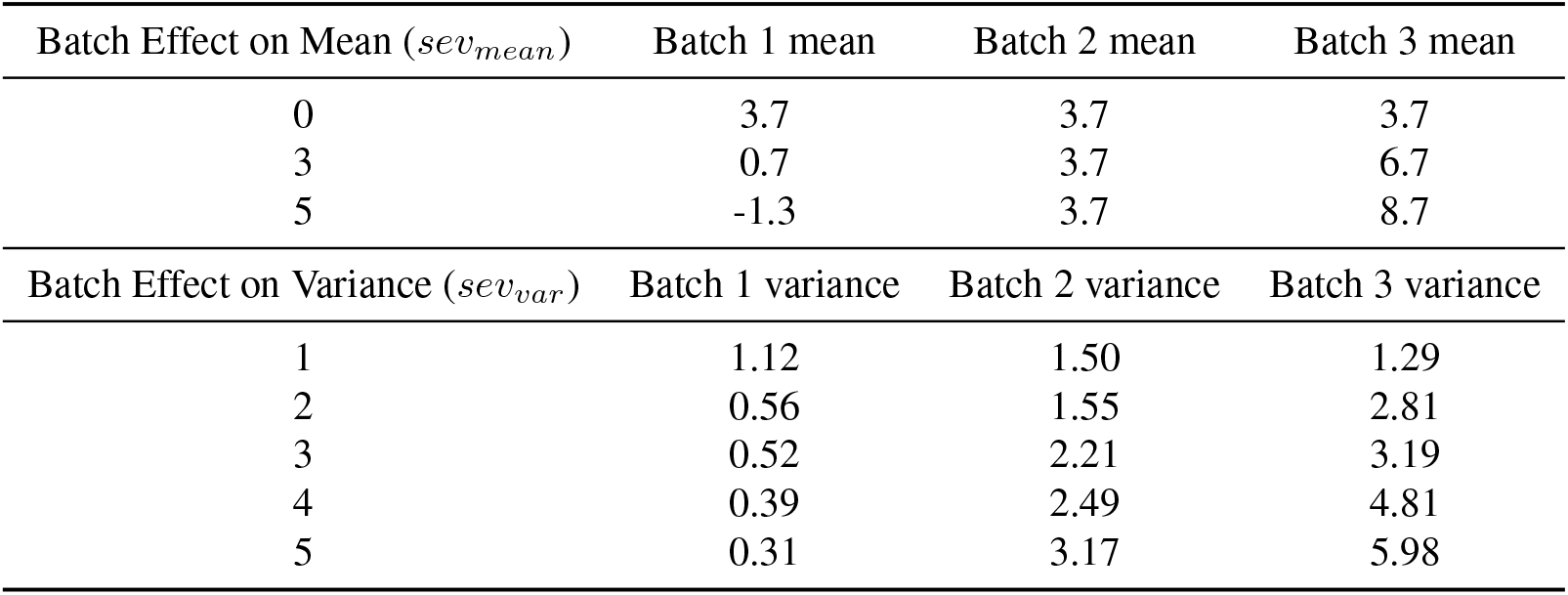
Levels of simulated batch effects in mean and variance of gene expression. We created mean shifts and fold changes (FC) in average variances of genes across the three simulated batches. ComBat assumes additive batch effect for the mean expression. For example, a batch effect on the mean of 3 means that we subtract 3 on average from expressions in batch 1, and add 3 on average to expression values in batch 3. Values in batch 2 are not altered. On the other hand, batch effect on the variance is multiplicative. So a batch effect of 4 on the variance means that in batch 1, the average gene variance is reduced to 1/4 of the original variance in data, while in batch 3, the average variance is inflated 4 times. Variance of batch 2 is changed to an intermediate level. The first column records the parameters we used in simulations, and correspond to the titles in Figure 2. The remaining three columns show the moments of distributions from example simulated data.

**Figure 1:**
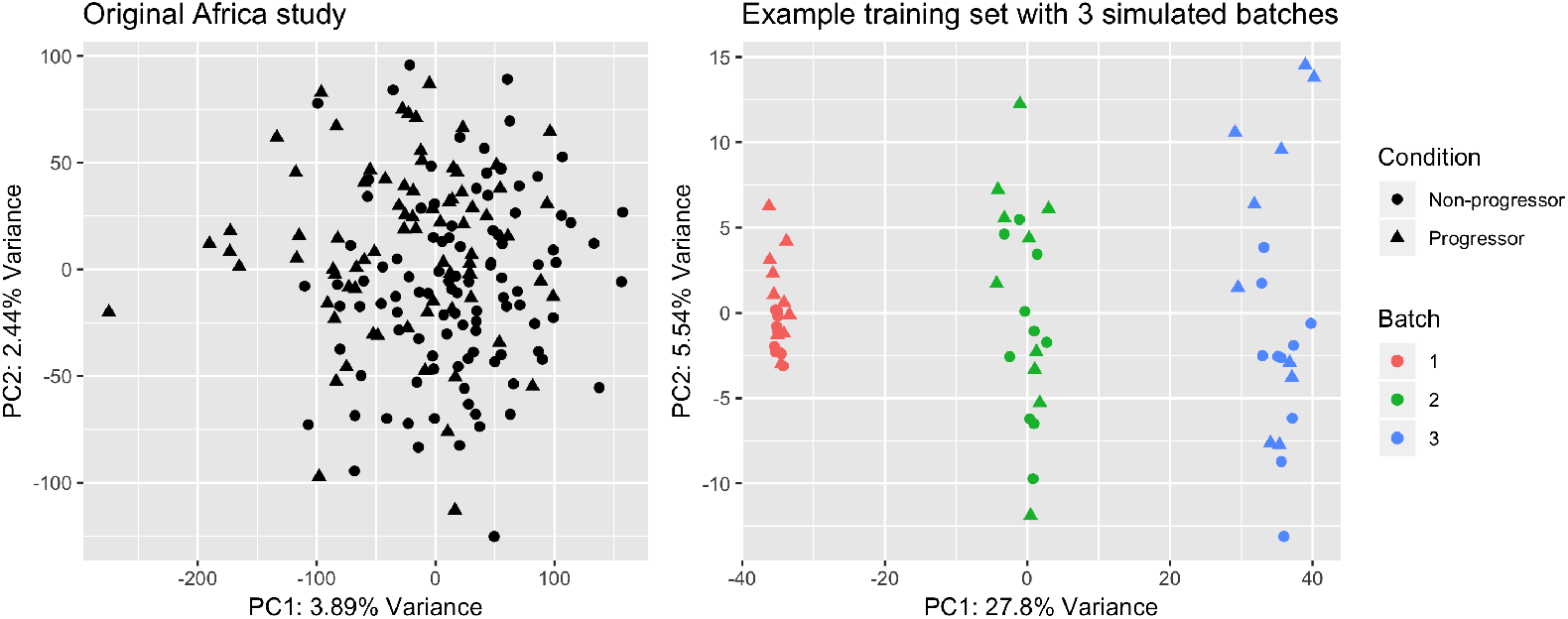
PCA of the original Africa study, and an example training set with simulated batch effect. We generated the training set by first taking 3 random subsets from the original data in the Africa study (Zak *et al.* (2016)), and treat them as 3 different batches. We then simulate batch effects in both mean and variance of gene expression across the three batches (see also Table 2). In this example, we simulated a mean difference of 3, and a variance difference of 4 fold across batches.

**Figure 2:**
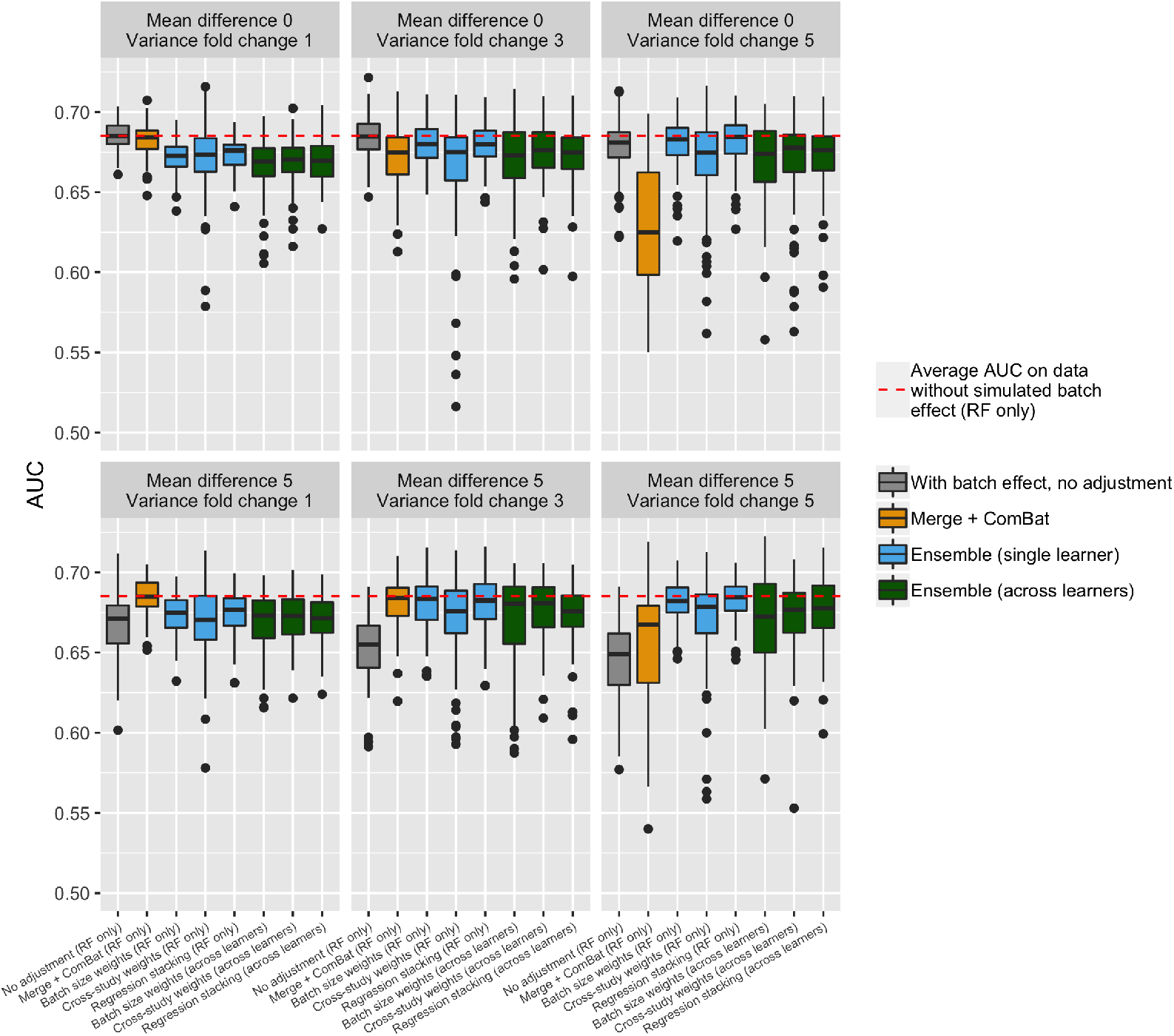
Comparison between ensembling and merging when using Random Forests. 3 out of our 5 choices of ensembling weights are displayed: batch size weights, cross-study weights, and stacking regression weights (See the Methods section for details). A full comparison using all 5 weighting methods, and a more refined grid of mean and variance differences, is in Supplementary Figure S3.

#### Comparing ensemble learning with merging after batch correction

We then use the dataset with simulated batch effects to train classifiers for predicting patient phenotypes. We perform the ensemble learning strategy as described above. For the merging strategy, we pooled the three batches together, and applied ComBat to remove batch effects. We then used the whole adjusted data to train a single model, and make one set of predictions on the independent test set. We trained learners LASSO (Tibshirani (1996)), Random Forest (RF, Breiman (2001)), and Support Vector Machines (SVM, Cortes and Vapnik (1995)), after performing the two batch adjustment strategies. In ensemble learning, we evaluated aggregating predictions both from a single learning algorithm (*L* = 1), and across all algorithms (*L* = 3). The accuracy of performance was measured by the area under ROC curve (AUC). We repeated batch correction and predictions to generate 100 discrimination scores, and compared them between the two strategies.

### Applying ensemble learning to address real data batch effects

To demonstrate a realistic application setting for our ensemble learning method, we took three studies from our TB cohort: Africa, US, and India. We iteratively treat each of these studies as the independent test set. The remaining two studies are used as the two batches forming the training set.

The original data contains more than 15000 genes, resulting in too unfavorable a situation so serve as a comparator for our methods. For example, LASSO was not able to get a prediction AUC above 0.5 on the independent samples. We thus prefiltered genes to select a subset of the 1000 most highly variable, and used the same subsets of genes for both ensembling and merging.

We trained 3 learning algorithms: LASSO, RF, and SVM, and integrated predictions from all three learning algorithms in the ensemble framework. The remaining methods for batch effect adjustment, predictions, and model evaluations are the same as those for simulation studies. We performed 100 bootstrap replicates on the test set, to obtain a confidence interval for model performance scores.

## Results

### Impact of mean and variance batch effects on discrimination of predictions

Figure 2 summarizes results over 100 simulated datasets, representative of the patterns we observe across simulation studies. The learner is Random Forests for both merging and ensembling. On the original training study without simulated batch effect, Random Forests achieve a 0.685 AUC on the test set. When adding batch effects to the data, we observed drops in discrimination in the test set. Mean and variance batch effects affect prediction performance in different ways. Model discrimination is not strongly affected by batch if batch effect only affect the variance. A sufficient size of the mean differences across batches in the training set is necessary to cause a drop in prediction accuracy. On the other hand, when batch affects the means, an increase in variance differences across batches will lead to a further drop in discrimination.

### Ensemble learning achieves higher discrimination than merging after a turning point on the severity of batch effects

Table 2 shows the levels of batch differences we created in simulation studies, corresponding to the results in Figure 2. We considered ensembling both using a single learning algorithm, and using all three learning algorithms together, and we observed similar results. At a low level of batch effects, with no mean difference and a variance fold change smaller than 3, the merging method yields better discrimination. However, we observed a turning point in the severity of batch effect, after which ensemble learning starts to achieve higher discrimination.

The turning point in the magnitude of batch effects differs by the selected learning algorithms. Supplementary Figures S2, S3 and S4 show the simulation results of training with all algorithms. When training with SVM, for example, we see that discrimination from ensemble learning is already comparable with that using the merging method at no mean batch effect and a variance fold change of 2, though at this level, merging still out-performs ensembling when training with Random Forests. Also, at high level of batch discrepancies, stacking regression weights yield better prediction results than the sample-size weights when building ensemble with SVM alone, though they are more comparable when using Random Forests.

We also observed that ensembling across different learning algorithms does not necessarily improve the final prediction compared to ensembling with a single algorithm. We see in Figure 2 that ensemble learning across learning algorithms generates worse performance than using only Random Forests. The optimal weighting approach also depends on the learners involved in the ensemble. When using Random Forests only, the sample-size weights and the stacked regression weights generate better accuracy than the cross-study weights. But the latter is better in integrating across all algorithms. Note that despite the difference in rankings of the three types of weights, all three ensemble weighting methods out-perform merging and batch correction with ComBat at high level of batch differences.

Finally, we repeated the simulations with a larger sample size, and larger number of batches. To increase batch size, we took 3 subsets as batches, each containing 20 non-progressors and 20 progressors, in contrast to our previous results which use 10 individuals per condition per batch. For a larger number of batches, we took 5 subsets of subjects as simulated batches. We observed consistent patterns in both situations, as shown in Figure 3 and Supplementary Figure S5.

**Figure 3:**
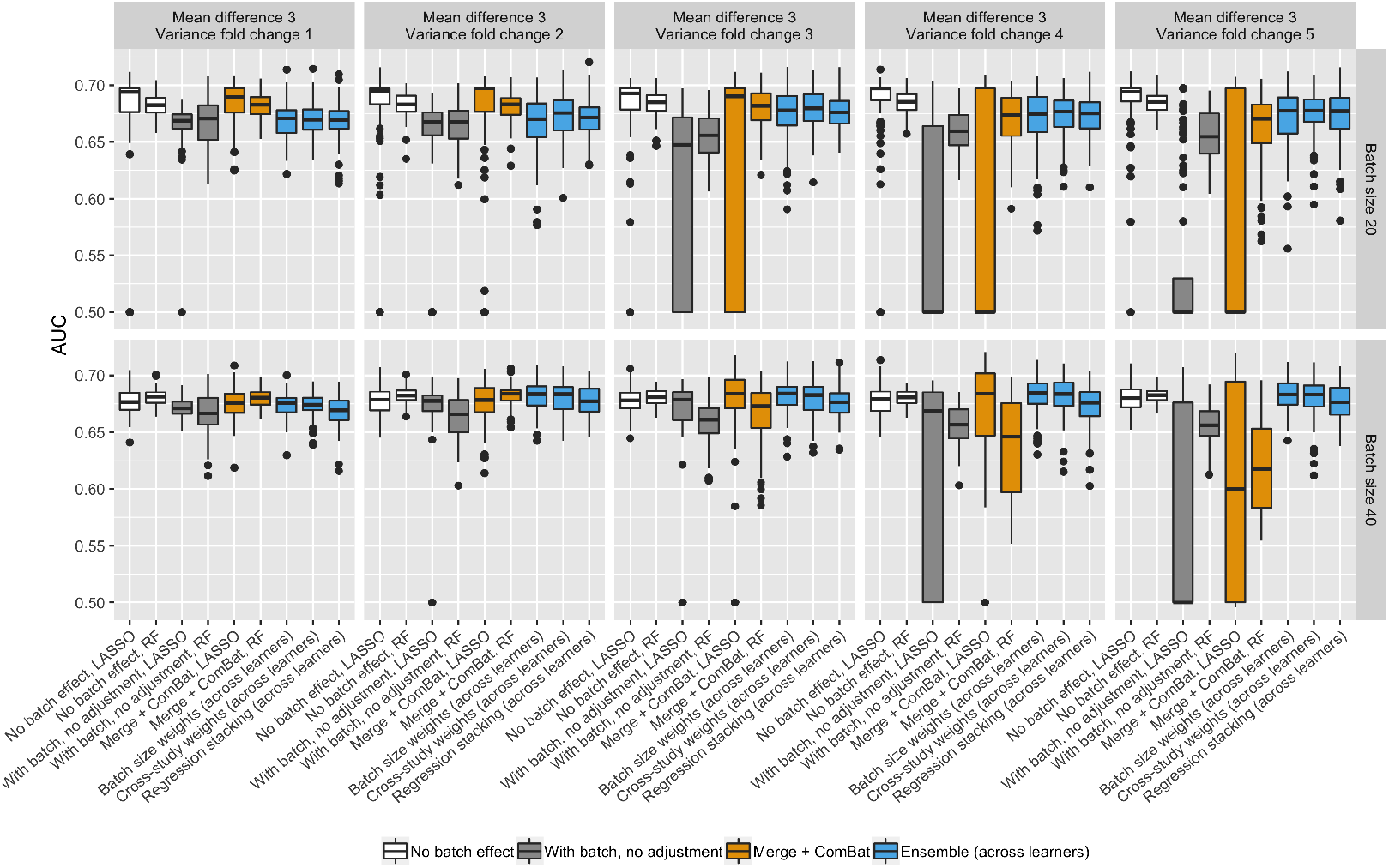
Comparison of results between batch sizes N=20 and N=40. In both cases, we have a completely balanced study design in each of the 3 batches. For N=20, we randomly sample 10 positive and 10 negative samples. For N=40, we sample 20 in each class. In both sample sizes, we observe that merging out-performs ensembling with smaller batch effects. However, as those increase, ensembling gains better discrimination.

### Application to predicting tuberculosis disease phenotype

We now present a classification case study using real data with batch effects. Specifically, we selected three batches from the TB c ohort, and iteratively treat one as test set, and use the other two as two batches for training. We applied both ensembling and merging to address batch effects, and trained 3 types of learning algorithms. Ensemble prediction are aggregated from all algorithms. To obtain a confidence interval of model performance, we generated 100 bootstrap samples from the test set.

Figure 4 shows the average AUC across the 100 bootstrap replicates, obtained from predictions in each of the three studies. The error bars in the figure display 95% confidence intervals. We observed that, except for the stacking weights when the US is the test set, the average AUC of ensembling using any of the three types of weights is better than that of merging in all studies. When using stacking weights and US as the test set, we found that ensemble assigns most of the weights to the SVM model trained from the Africa study (Supplementary Figure S6). This model achieves accurate discrimination in both Africa and India, the two batches for training, while inferior preference in US compared to the other models (Supplementary Table S1). As a result, stacking weights generate less favorable predictions than cross-study weights, where weights are more evenly spread across the single learners. Cross-study weights only consider prediction performance across studies, while stacking weights, at least as implemented here, consider both within and across study performance. With only two studies, the importance given by stacking weights to cross-study performance is at its minimum. As a result this method is more vulnerable to overfitting than cross-study weighting.

**Figure 4:**
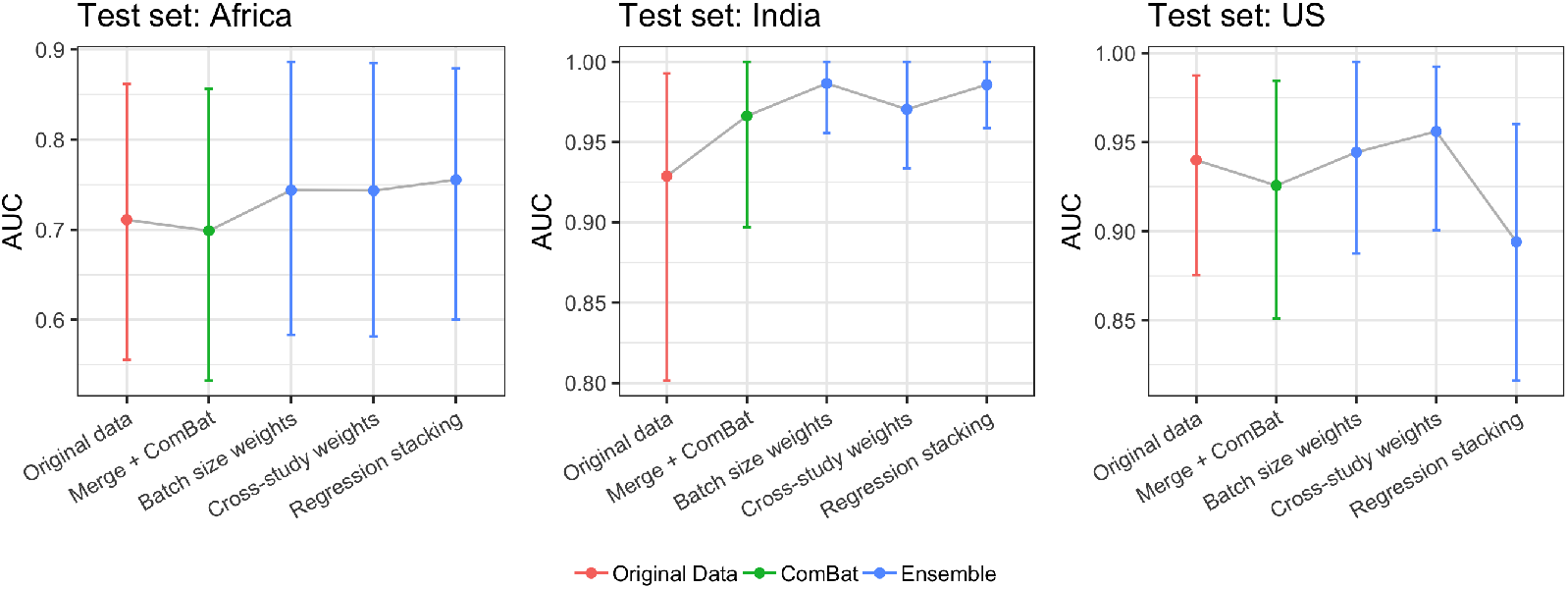
Application of ensemble learning to predicting active TB against latent infection. We iteratively select one of the three studies in Table 1 as the independent test study. The remaining two studies are viewed as two “batches” in the training set. The two batches in this setting represent a strong level of batch effects. We trained LASSO, Random Forest, and SVM, then aggregated predictions from all three algorithms to construct the ensemble. The figure shows average prediction performance over 100 bootstrap samples of the test data, with error bars showing 95% confidence intervals of performance measures. Except when using stacking weights to predict on the US study, the average performance using the three ensemble strategies are better than the merging strategy. Comparisons within each bootstrap sample are summarized in Table 3.

**Table 3:** Percentages of bootstrap test samples where ensembling yields strictly higher AUC than merging and batch adjustment. Rows indicate validation datasets (see caption to Figure 4) and columns indicate ensembling methods, each compared to merging.

| Test set | Batch size weights | Cross-study weights | Stacking regression weights |
| --- | --- | --- | --- |
| Africa | 97% | 91% | 94% |
| India | 84% | 48% | 76% |
| US | 73% | 85% | 20% |

Due to the relatively small sample size in the US and India studies, we also observed a high variance in model performance, especially when Africa is used as the test set. Therefore, we also compared ensembling against merging within each bootstrap sample. The proportion of bootstrap samples where ensembling generates strictly higher AUC than merging are summarized in Table 3, which shows that in all three test sets, addressing batch effects via ensemble learning yields better-performing models with respect to discrimination. Batch-size weights, while not explicitly trained to reward cross-study replicability, are more consistently outperforming merging than other weights in this dataset, likely because of the challenges in learning weights with a small number of batches. These results are broadly consistent with our observations in simulations that when there are severe differences across batches, ensemble learning is the better strategy to address the impact of batch effects on prediction.

## Discussion and Conclusions

We proposed a novel perspective for addressing batch effects when developing genomic classifiers. Our proposal is to use multi-study ensemble learning, treating batches as separate studies. We provided both realistic simulations and real data application examples to compare the ensembling method with the traditional approach, which is to analyze all batched together to remove batch effects, and use the adjusted data for prediction. We observed in both simulations and real data that, though merging is able to generate better performing models when batch effects are modest, ensemble learning achieves better discrimination in independent validation after a certain level of severity of batch effects. We explored different training algorithms, different batch sizes and number of batches, and observed consistent patterns of such transition. The specific level of batch severity where the transition happens differs by the algorithm. These observations are consistent with those described in Guan *et al.* (2019), which provides theoretical insights into the comparison between merging and ensembling in training cross-study learners.

The philosophy behind the standard approach of merging and batch adjustment is to remove the undesired batch-associated variation from as many of the genomic features as feasible, and then use the “cleaned” data in classification as though the batch effects never existed. This has been the standard in the literature and can be quite successful (Riester *et al.* (2014); Luo *et al.* (2010); Engchuan *et al.* (2016)). Multi-study ensemble learning provides a different perspective: ensemble weights reward prediction functions that, while trained in one batch, continue to predict well in other batches. These are likely to avoid using features affected by batch effect, in contrast to cleaning them.

Our results emphasize the importance of understanding the heterogeneity introduced by batches before developing a classifier. Since the approach which results in better prediction accuracy changes by the level of heterogeneity, we suggest careful diagnostics of the level of heterogeneity in the dataset, before choosing the learning approach. There are established methods to detect the degree of batch effects in data. For example, BatchQC (Manimaran *et al.* (2016)) offers sample and gene-wise statistical tests to explore mean, variance, as well as higher-order moments in batch distributions, and provides useful visualization tools to describe the severity of batch effects in data.

In simulations, we compared ensembles based on a single learning algorithm, to one based on multiple algorithms. Though the latter represents the common perspective of ensemble in practice, we found that integrating across multiple learning algorithms does not necessarily improve prediction performances than using a single algorithm. We also compared five kinds of weighting strategies to integrate the predictions. The ranking of performance using different weighting strategies also depends on the specific dataset and algorithm. Additionally, there could be ways to further improve the ensemble performance by developing other weighting methods that are not considered in this study. All ensembles considered are very small, because the number of batches *B* and the number or learners *L* are both small. Recently Ramchandran *et al.* (2019) compared an ensemble strategy based on RF to one based on cross-study weighting of the component trees in the RF, showing improvements from direct reweighting of trees. This method may also prove effective in our context, as individual trees are more parsimonious than the whole forest, and may more effectively avoid features affected by batches.

Our study has several limitations. First, the ensemble learning approach requires that each batch contains sufficient samples to train a prediction function. This assumption may limit our approach to sufficiently large datasets. However, most if not all batch effect adjustment strategies require a reasonable number of samples in each batch to accurately and robustly estimate batch effects. Having limited number of samples in a batch will negatively affect not only our proposed methods, but also the traditional methods based on merging. We speculate that methods like ComBat might, however, be able to effectively operate with smaller batches than ensembling.

We focused on using ComBat for batch effect adjustment after merging. ComBat is not the only option, but remains one of the most popular batch effect adjustment methods, especially in the case with known sources of batch effect. Our simulations of batch effects are all based on the generative model of ComBat, which, while plausible, is one among many possible models. Using the same batch effect model for data generation and analysis provides a lower bound to the effectiveness of our proposal, as any other data generating approach would be less favorable to ComBat than the one we used. In data from other generating mechanisms, the advantages of ensembling should be more pronounced, and may set in at lower levels of batch effects.

We provided a realistic application example using the tuberculosis studies. Restricted by the study design of this collection of studies, we showed an example with 2 batches, and relatively small batch sizes in two batches. It would be interesting to further explore our approach on data with larger batch sizes, or a larger number of batches. Ours are somewhat extreme conditions for this type of analysis. Larger batches facilitate both the estimation of batch effects and the training of batch-specific predictors. A larger number of batches facilitates learning about the higher level distributions in ComBat, and would afford ensemling a better opportunity to find stable signal across a larger number of batches, a strength of the method that is not highlighted here.

Related, treating two studies as batches mimics a high level of batch differences, for the discrepancy between the two “batches” in this setting includes both biological and technical differences. Thus we include more sources of heterogeneity than normally considered as batch effect. Specifically, a batch effect is usually defined as variations originated from technical differences across repeated experiments performed on the same platform, such as differences in lab environment, protocols or reagents. In our application example, however, each batch targets a different population, which means there likely exists additional genetic variations across the groups. Also, the three studies are generated on different platforms. The African cohort was sequenced on Illumina HiSeq 2000, the Indian study was produced on Illumina NextSeq 500, and the US study used microarray (Affymetrix Human Gene 1.1 ST Array). Differences across these technologies are also beyond what is typically considered as batch effects. Despite that, we believe this example is still helpful to illustrate our methodology, and our observations offer valuable practical guidelines in addressing batch effects in genomic classifier development.

### Reproducibility

Code to reproduce the results in this paper is available at https://github.com/zhangyuqing/bea_ensemble.

## Acknowledgement

This work is supported by grants NSF-DMS 1810829 and NIH-NCI 4P30CA006516-51 (Giovanni Parmigiani) and 5R01GM127430-02 (W. Evan Johnson).

