## Supplementary Materials for "Robustifying genomic classifiers to batch effects via ensemble learning"

*: Joint last authors

#### 1 Ensemble weighting methods

The weights are defined as described in Patil and Parmigiani [2018]. Here we use the same notations as in the Method section in the main article. Specifically, we fit each of $L$ learning algorithms to each of the $B$ batches in the training set. Models obtained by fitting algorithm $k$ on batch $b$ is denoted as $\hat{Y}_b^k(\mathbf{x})$, and their corresponding weight is $w_{kb}$. We compare five kinds of weights:

- Simple average. Weights for all models are constant $w_{kb} = 1/LB$.
- Batch-size weighted average. $w_{kb} = n_b/(L\sum_r n_r))$, where $n_r$ is the sample size of batch $r$. Under these weights, the larger batches get higher weights.
- Cross-study weights. We use each model $\hat{Y}_b^k(\mathbf{x})$ to make predictions in each of the other batches aside from $b$. In doing so, we construct a matrix of model performance $Z^k$ for each algorithm $k$, where the element at $(i,j), i \neq j$ is $z_{ij}^k = \text{MXE}_{ij}^k = \frac{1}{n_j}\sum(y_j \log \hat{Y}_i^k(\mathbf{x_j}) + (1-y_j)\log(1-\hat{Y}_i^k(\mathbf{x_j}))$, which refers to the mean cross-entropy loss of predictions on batch $j$, using model $\hat{Y}_i^k$. We then summarize these matrices across the batches for prediction $(j)$, to obtain the unscaled weight for model $\hat{Y}_b^k$: $z_{kb} = (\frac{1}{B-1}\sum_{j,j\neq b} z_{bj}^k)^{\frac{1}{2}}$. Finally, the scaled weights $w_{kb}$ is defined as

$$w_{kb} = \frac{1}{C}|z_{kb} - \max_{k',b'}(z_{k'b'})| \tag{1}$$

  where $C$ is a normalization factor which ensures that the weights have a sum of 1. This approach assigns zero weight to the model with the worst average performances on the rest of the training batches (in other words, worst generalization performances estimated using the remaining batches within the training set).

- Aggregated regression weights. We use each model to make predictions in each other batch. As a simplified notation, we use $p_{ij}^k = \hat{Y}_i^k(\mathbf{x_j})$ to describe the predicted probabilities on batch $j$ using model $\hat{Y}_i^k$. Note that $j$ can equal $i$ in this case. We calculate the coefficients of non-negative least square regression for each batch $j$:

$$\beta_j^k = (\beta_{1j}^k, \beta_{2j}^k ..., \beta_{Bj}^k) : y_j \sim p_j^k \quad (2)$$

where $p_j^k = (p_{1j}^k, p_{2j}^k, ..., p_{Bj}^k)$ is a matrix of predicted probabilities, with each column being predictions using model trained from one batch. The weights are then defined as the batch-size weighted average of vector $\beta_j^k$:

$$(w_{k1}, w_{k2}, ..., w_{kB}) = \sum_j n_j * \beta_j^k / \sum_j n_j \quad (3)$$

or equivalently, $w_{kb} = \sum_j n_j \beta_{bj}^k / \sum_j n_j$.

- Stacking regression weights. Stacking weights are constructed in similar ways as the aggregated weights, only that now, we stack the labels and predictions for all training batches. We use $Y = (y_1^T, y_2^T, ...y_B^T)^T$, and $P = (P_1^k, P_2^k, ...P_B^k)$ to denote the stacked labels from all training batches and predictions in these batches. $P_b^k = ({p_{b1}^k}^T, {p_{b2}^k}^T, ..., {p_{bB}^k}^T)^T$ is the stacked predictions on every batch, using model $\hat{Y}_b^k$ from batch $b$. The weights are then computed through a single non-negative least square regression:

$$(w_{k1}, w_{k2} ..., w_{kB}) : Y \sim P \quad (4)$$

We selected the batch-size weighted average, cross-study weights, and stacking regression weights as representatives to show in the main paper. Results using all five weighting methods are summarized in Supplementary Figures S2, S3, and S4.

#### 2 A schema of our approach

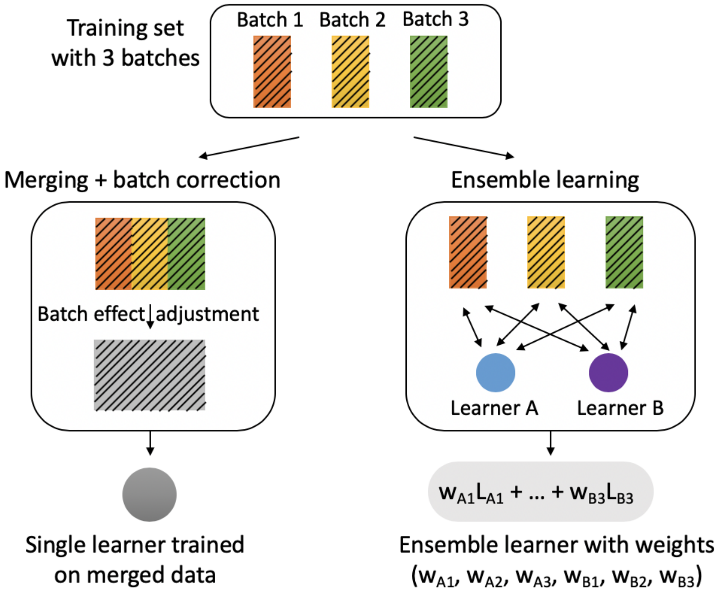


Figure S1: **A schema comparing batch effect adjustment via ensemble learning and merging.**

#### 3 Additional simulation results

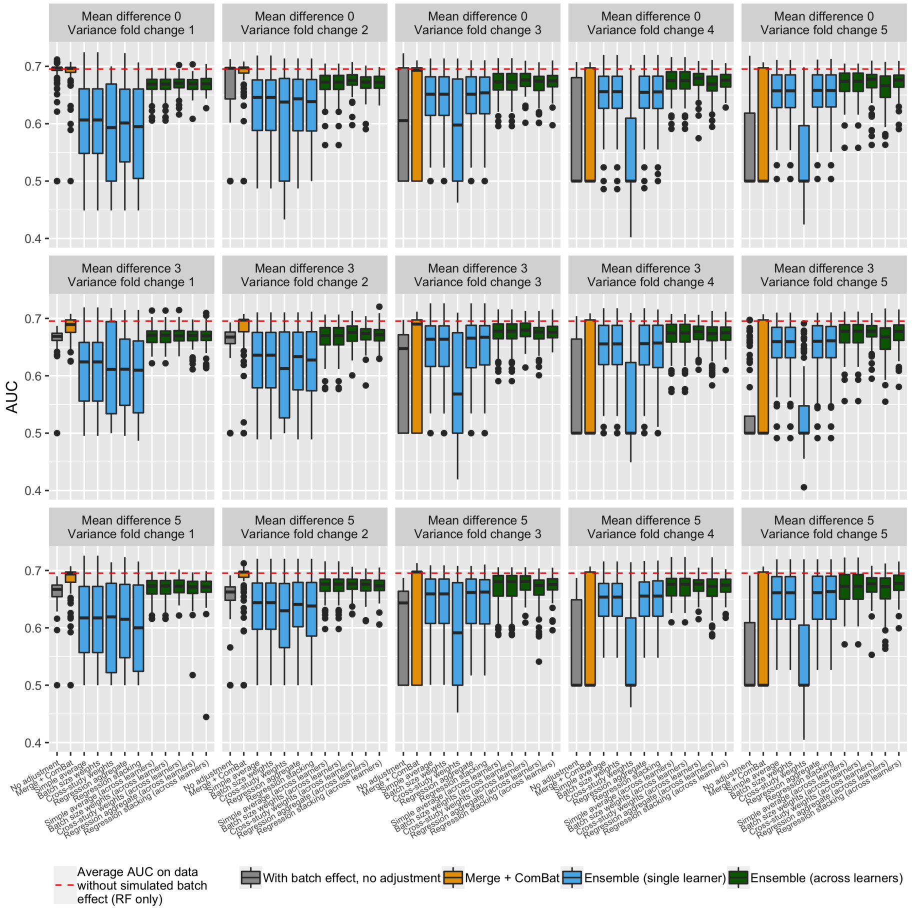


Figure S2: **Full comparison between learner and data integration - LASSO.**

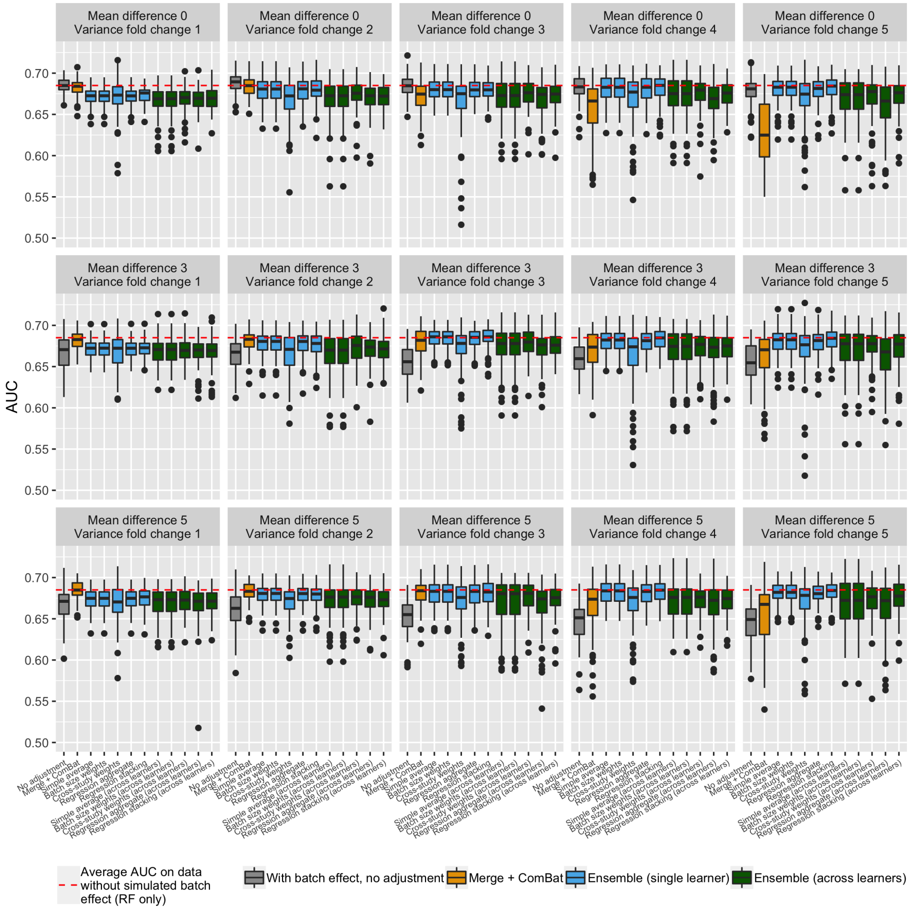


Figure S3: **Full comparison between learner and data integration - Random Forest.**

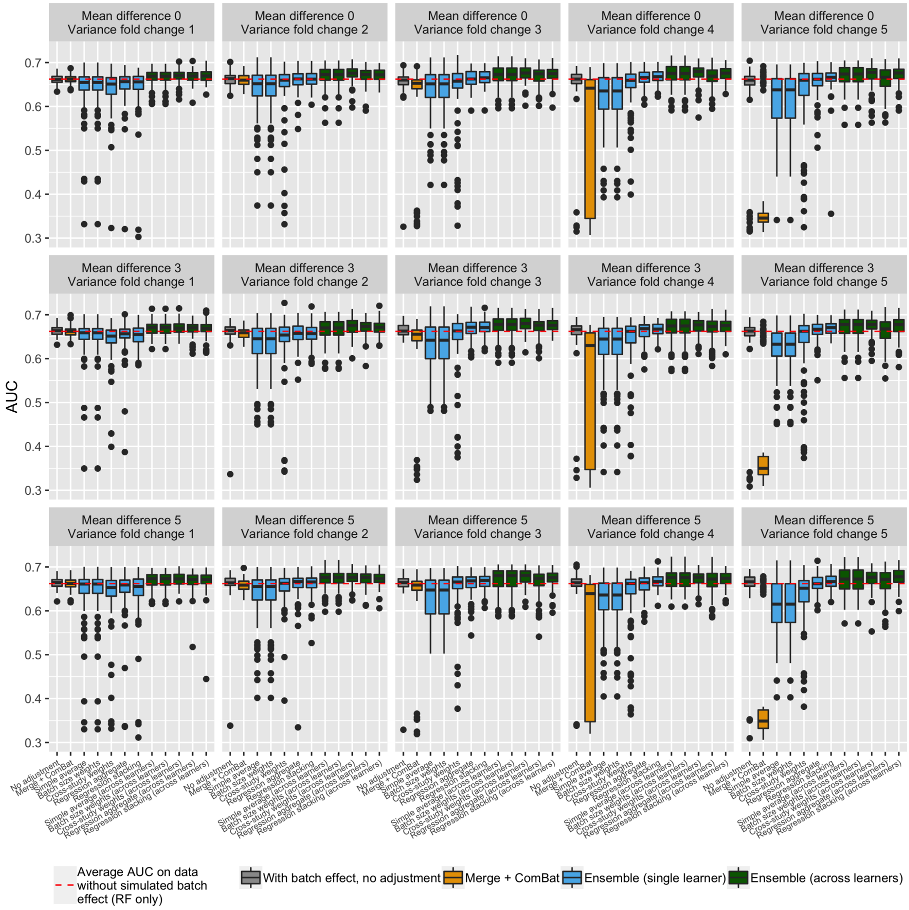


Figure S4: **Full comparison between learner and data integration - Support Vector Machines.**

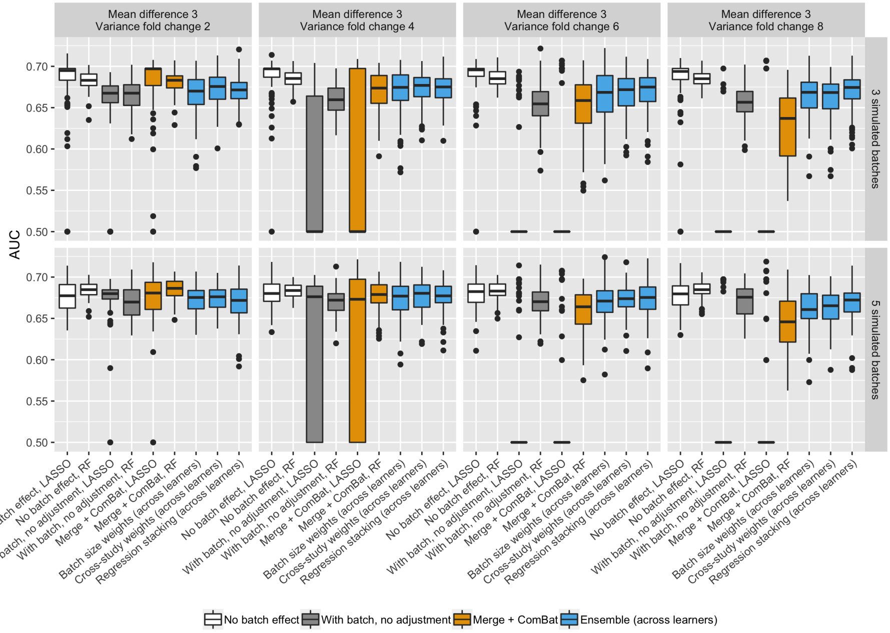


Figure S5: **We observed consistent results between simulating 3 and 5 batches.** In both cases, we have a completely balanced study design in each of the batches. Same as in the other simulations, we found that merging outperforms ensembling at a lower level of batch effect. While as the severity of batch effect increases, ensemble starts to gain better prediction abilities.

### 4 Additional real data results

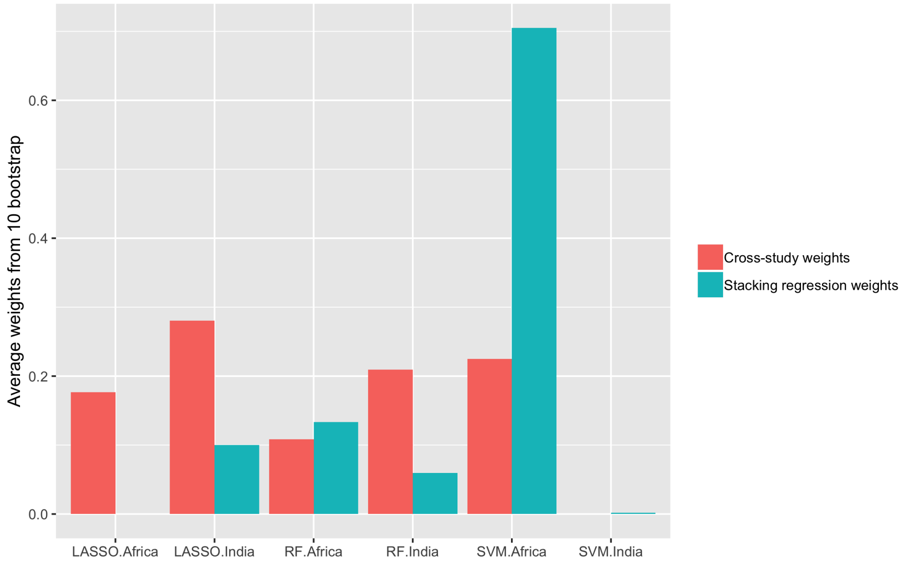


Figure S6: **Comparison of cross-study weights and stacked regression weights in the real data example, when US is the test set.** The barplot shows the average weights on each model across 10 bootstrap samples.

We observed that merging out-performs ensembling only when using the stacking regression weights and when making predictions on the US study. To provide further insight, we compared cross-study weights and stacked regression weights from 10 bootstrap samples. When using stacked weights, ensembling assigns most of the weights to SVM trained from the Africa study, which has good performance in Africa and India, but inferior performance in the US. In comparison, cross-study weights are more evenly spread across single learners.

| Learner | Training | Batch1 AUC (Africa) | Batch2 AUC (India) | Test AUC (US) |
|---|---|---|---|---|
| LASSO | Africa | 0.88 | 0.98 | 0.96 |
| LASSO | India | 0.83 | 1.00 | 0.95 |
| RF | Africa | 1.00 | 0.91 | 0.79 |
| RF | India | 0.71 | 1.00 | 0.99 |
| SVM | Africa | 1.00 | 0.91 | 0.81 |
| SVM | India | 0.74 | 1.00 | 0.98 |

Table S1: **Summary of single learner performances from one example boostrap, when using US as the test set**. Stacked regression weights are mainly assigned to SVM model trained in Africa, which has over 0.9 AUC in both batches, Africa and India, of the training set. However, its test set performance is less ideal compared to the other models, resulting in worse discrimination performance than merging.
